# Automated Gene Data Integration with Databio

**DOI:** 10.1101/768077

**Authors:** Robert W Reid, Jacob W Ferrier, Jeremy J Jay

**Affiliations:** Department of Bioinformatics and Genomics, College of Computing and Informatics, University of North Carolina at Charlotte, Charlotte, North Carolina, USA; North Carolina Research Campus, Kannapolis, North Carolina, USA

## Abstract

**Summary:** Databio is capable of providing fast and accurate annotation of gene-oriented data sets, coupled with an integrated identifier conversion service to empower downstream data mining and computational analysis. Databio is enabled by fast real-time data structures applied to over 137 million unique identifiers, and uses automated heuristics to permit accurate data provenance without highly specialized knowledge and bioinformatics training.

**Availability and Implementation:** Freely available on the web at https://datab.io/. Source code and binaries are freely available for download at https://github.com/joiningdata/databio/, implemented in Go and supported on Linux, Windows, and macOS.

## 1 Introduction

Although sequencing and other high-throughput data production technologies are increasingly affordable, data analysis remains a significant factor in the cost of -omics studies (Mardis, 2010). Without improving the ability to automate data integration and interoperation, the cost of analysis will continue to impede access to precision medicine for underserved populations with limited resources. Many tools and standards have been developed around the concept of a central “Data Commons”, but the path forward remains unclear (NIH, 2019), and current large-scale data repositories continue to be highly specialized and difficult to apply in generalized study. Despite the acceptance and proliferation the FAIR data guidelines (Wilkinson et al., 2016), current data provider implementations focus on descriptive metadata and keyword-oriented search applications, leaving the majority of detailed gene and other -omics data inaccessible and difficult to discover through computational means.

Data producers recognize the need to enable greater access to hosted data, but there are no well-accepted machine-readable means for annotating the contents of data sets across the biomedical landscape (National Research Council (US) Board on Biology et al., 2000). The lack of available standards and tools make it a cumbersome and time-consuming task to properly annotate identifier sources, record their provenance throughout an analytical process, and track subsequent data quality metrics. As a result, the majority of useful scientific results remain buried in supplementary tables, figures, and poorly indexed data archives. Efforts to exploit these types of results often require specialized knowledge, thus there is a need to simplify the discovery and retrieval process.

We present Databio, a novel framework for automating the extraction, annotation, and integration of gene-oriented data sets. Databio automates data parsing and identifier detection, and streamlines many common tasks to provide a point-and-click approach to data manipulation and integration across a broad spectrum of applications in life sciences research and translational medicine. This ability to quickly and accurately streamline complex tasks will enable faster and better analysis of -omics data.

## 2 Implementation and Available Data

Databio is implemented as a web-based data portal (https://datab.io) that allows users to interact with the embedded tools using an interactive web browser-based interface.

User data uploads are first handled via an automatic detection framework that determines the source data format (see top Fig. 1). The current implementation supports Tab-separated values (TSV), Comma-separated values (CSV), and Excel 2007+ spreadsheets (XLSX). Records(rows) and fields(columns) within these documents are exposed to the rest of the application through a modular interface allowing for support for more data formats in future software updates. Heuristic techniques are applied to the parsed data to remove headers and determine field labels, allowing for a more descriptive display interface (see Fig. 1).

**Figure 1.**
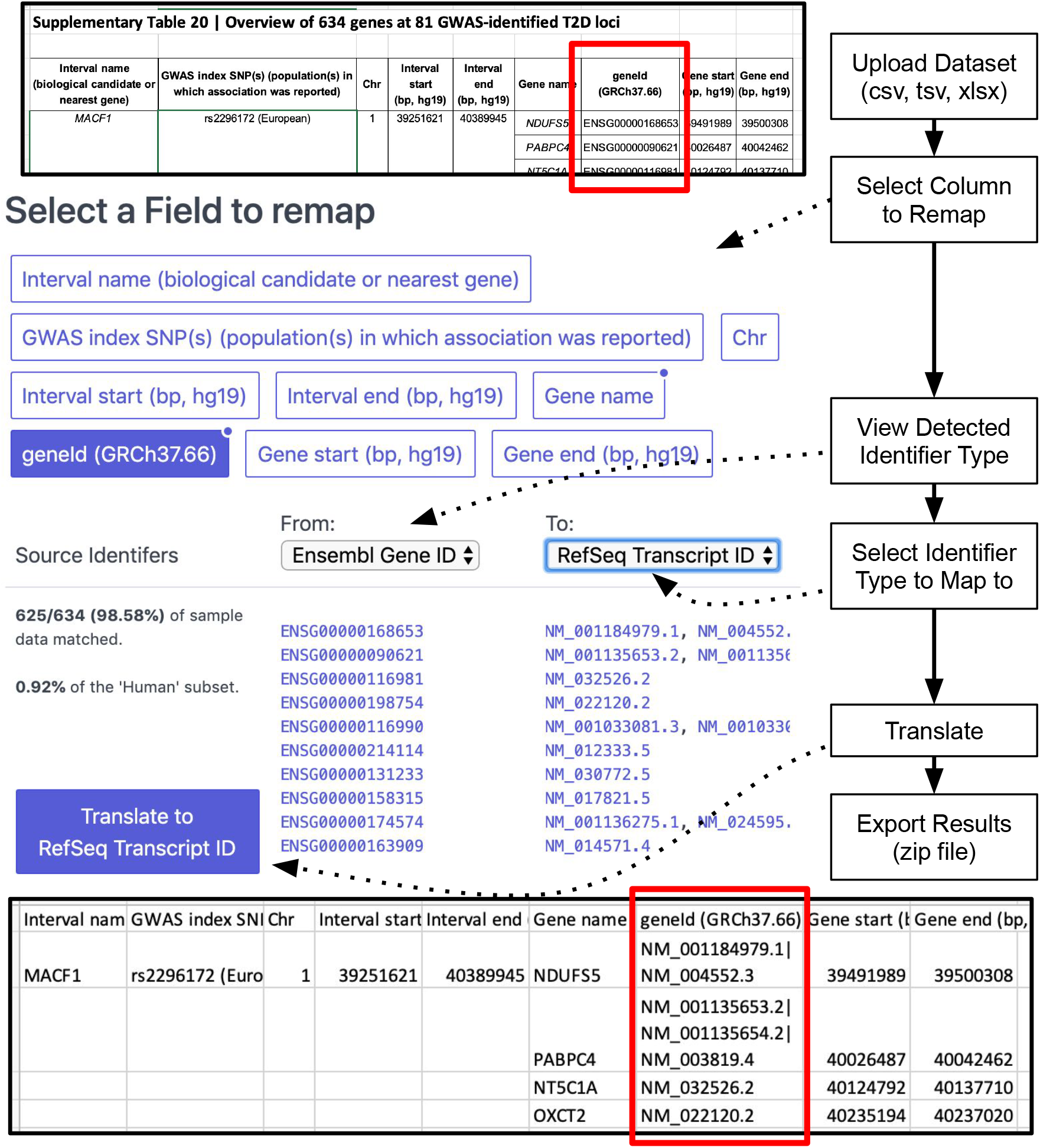
Databio web interface workflow showing data upload (including Excel formatting, headers, and merged fields). Point-and-click field mapping allows selection of source and replacement gene identifiers. Results are then exported with new identifiers. Statistics, bibliography, and provenance files are included but not shown in figure.

Once fields are parsed, values are aggregated together and searched against our warehouse of multiple gene identifier data sources. Our current snapshot contains over 137 million unique gene, transcript, and protein identifiers and 92 million unique mapping pairs (Table 1). Despite the extreme scale of determining identifier source, this classification can be completed accurately in real-time (less than one second) using using Bloom filters for fast approximate matching (Bloom, 1970). The top hits for each field are collected (along with sample values) and returned to the web interface so that users can verify the accuracy of the predicted identifier type.

**Table 1.** Gene identifier sources loaded into Databio as of 2019-09-12.

| Name | Subsets | Total Unique | Reference |
| --- | --- | --- | --- |
| NCBI Entrez Gene | 39 | 25,295,958 | (Maglott et al., 2007) |
| RefSeq Transcripts | 1 | 2,211,841 | (Pruitt et al., 2005) |
| RefSeq Proteins | 1 | 40,574,328 | " |
| Ensembl Gene | 207 | 5,442,203 | (Zerbino et al., 2018) |
| Ensembl Transcripts | 207 | 9,000,822 | " |
| Ensembl Proteins | 207 | 6,923,465 | " |
| KEGG Genes | 6128 | 29,541,384 | (Kanehisa et al., 2008) |
| UniprotKB/Swiss-Prot | 1 | 18,493,595 | (UniProt Consortium, 2015) |
| HGNC Gene IDs | 1 | 42,050 | (Yates et al., 2017) |
| HGNC Symbol | 1 | 42,050 | " |
| HGNC Gene Names | 1 | 42,050 | " |
| OMIM Genes | 1 | 16,197 | (Hamosh et al., 2005) |

In addition to the classification index representation, the Databio database also contains mappings that allow supported identifiers to be translated into other identifier types. Although this common task has been supported by other tools such as David, Uniprot, and BioMart (Dennis et al., 2003; UniProt Consortium, 2015; Smedley et al., 2015), these tools require manual data manipulation, specialized knowledge of identifier sources, and cannot replace identifiers within the context of the original data file. Databio is able to translate identifiers in-place, removing multiple opportunities for error and keeping the data in context. These changes are applied to the existing data schema and exported to a CSV-format data set that can be readily imported into other tools for subsequent analysis (see bottom of Fig. 1).

Further easing the burden of data manipulation on the user, Databio is able to track important data quality issues such as missing identifiers and ambiguous mappings. The Databio warehouse maintains a record of publication and citation info for each identifier source, the last fetch and access dates, and analysis logs describing processing steps and data quality metrics. Using this information, Databio can establish that necessary metadata for publication, distribution, and reuse is present and accurately tracked. This ensures that data consumers know the state of a data set including access dates, citations, and relevant usage limitations.

## 3 Usage

For example, a study identified 634 genes associated with Type 2 Diabetes GWAS loci (Fuchsberger et al., 2016), and provided the results in a Supplementary Table (see top, Fig. 1). We want to look for relationships between the RefSeq Transcript sequences of the genes and the listed loci. However, searching for ‘ENSG00000168653’ in RefSeq currently yields no results, and the gene Symbol ‘NDUFS5’ returns 19 Human results. One must translate the gene identifiers into more specific RefSeq Transcript IDs.

Upon visiting the Databio site, the user is able to upload this Excel file (or any other TSV, CSV or XLSX data file) even though it does not fit a pre-determined field layout. Column names (fields) are automatically parsed and identified for selection on the second page (see top, Fig. 1). Fields with high-quality automated classification are marked with a circle in the top right corner to indicate a high correspondence to a known Databio identifier source (For example, the blue box “geneId (GRCh37.66)” in Fig. 1). The user is then able to click on the field name that they want to remap. The exact match rate, as well as the percent coverage of the corresponding source dataset, is shown to the user under the ‘Source Identifiers’ header on the left.

We can see that for this example, even though the file did not explicitly mention the source of gene identifiers, Databio easily determined them to be Ensembl Gene IDs. For other data sets, if there is more ambiguity to the identifiers (e.g. integers), the user can use the drop-down on the left to see the other matched identifiers sources and find the most appropriate choice. The user can then choose the desired identifier type to map to, using the drop-down on the right, and an automatically generated list of identifiers that map to the original identifier source. Changing either the ‘to’ or ‘from’ drop-down selections automatically updates to display a sample of the original identifiers from the uploaded data, and the associated remapped identifiers so that the user can confirm expectations. Finally, the user may begin the translation processing, which leads to a new page including the remapped data file for download, statistics, some text describing the methods and data sources used with a bibliography and analysis logs. This information is all available in a compressed ZIP archive ensuring that important information is delivered together as one unit.

## 4 Conclusion

Databio automates and streamlines the process of gene identifier translation, enabling new approaches to data-driven discovery by lowering the burden of data manipulation and prior knowledge of biomedical resources. Support for more identifier sources, more data formats, and chained identifier conversions (A → B → C) will greatly increase the utility of Databio across the life sciences. In addition, future computational analyses will build upon this base, enabling data set search based on related data contents and not just shared metadata. Together these improvements will enable future machine learning applications by removing the need for manual intervention in data import processes, shortening learning times and improving the pace of data-driven discovery.

